# Extraction and selection of high-molecular-weight DNA for long-read sequencing from *Chlamydomonas reinhardtii*

**DOI:** 10.1101/2022.08.16.504088

**Authors:** Frédéric Chaux-Jukic, Nicolas Agier, Stephan Eberhard, Zhou Xu

## Abstract

Recent advances in long-read sequencing technologies have enabled the complete assembly of eukaryotic genomes from telomere to telomere by allowing repeated regions to be fully sequenced and assembled, thus filling the gaps left by previous short-read sequencing methods. Furthermore, long-read sequencing can also help characterizing structural variants, with applications in the fields of genome evolution or cancer genomics. For many organisms, the main bottleneck to sequence long reads remains the lack of robust methods to obtain high-molecular-weight (HMW) DNA. For this purpose, we developed an optimized protocol to extract DNA suitable for long-read sequencing from the unicellular green alga *Chlamydomonas reinhardtii*, based on CTAB/phenol extraction followed by a size selection step for long DNA molecules. We provide validation results for the extraction protocol, as well as statistics obtained with Oxford Nanopore Technologies sequencing.

## Introduction

In recent years, long-read sequencing technologies, such as the ones developed by Pacific Biosciences (PacBio) and Oxford Nanopore Technologies (Nanopore), have emerged as a solution to the pitfalls of short-read technologies in the detection of structural variants and in assembling repeated sequences and other complex regions (1). Additionally, because native DNA is used, long-read technologies can directly detect a variety of modified bases, including the most commonly studied methylated cytosines (2, 3). For their applications in genome assembly and structural variant detection, these technologies typically sequence DNA molecules ranging in size from kilobases to hundreds of kilobases as a continuous read. Reads traversing repeated sequences are necessary to correctly assemble neighboring regions, with longer reads enabling more contiguous genome assemblies. Today, the major bottleneck to sequence long reads comes from the ability to extract high-quality DNA devoid of polyphenol and polysaccharide contaminants with sizes compatible with this purpose. This is especially true for most plant tissues and algae cells, because polyphenols and polysaccharides are often co-extracted with DNA and can inhibit downstream applications such as sequencing (4, 5).

*Chlamydomonas reinhardtii* is a unicellular green alga that is widely used as a model organism to study photosynthesis and cellular motility (6), and is an organism of choice for biotechnological application, with many synthetic biology tools being currently developed (7, 8). In *C. reinhardtii*, as for other plants and algae, contending with phenolic and polysaccharide contaminants while preserving HMW DNA is a major challenge and requires an optimized protocol. PacBio and Nanopore sequencing have been performed on this organism, contributing to important advances in our understanding of its genome structure and content, base modifications and evolution (9–13). However, previous protocols did not include a size selection step, which can substantially enrich for longer molecules.

Here, we present a detailed protocol dedicated to efficiently extract and select HMW DNA from *C. reinhardtii* cells. The protocol minimizes DNA-shearing manipulations and comprises an additional step to enrich for HMW DNA. We validated the method by pulse-field gel electrophoresis (PFGE) and measurement of read length from Nanopore sequencing.

## Materials and Methods

The protocol described here is also published on protocols.io, dx.doi.org/10.17504/protocols.io.8epv59j9jg1b/v2 and is included as supporting information file 1 with this article.

### Nanopore sequencing

Sequencing libraries were prepared as per manufacturer’s recommendations, using NEBNext companion module (E7180S, NEB) and Ligation Sequencing Kit SQK LSK-109 (Nanoporetech), except for the ligation time, which we increased to 30 min. For each run, 500 ng were loaded on MinION flow cells (R9.4.1, Nanoporetech) and sequenced for 6h to 16h, depending on flow-cell kinetics. Libraries were loaded at least twice, with 1h wash using the manufacturer’s washing buffer (EXP-WSH004) between runs. Basecalling was performed using Guppy (version 4.3.4) with parameters set to “high accuracy”.

## Results

We extracted genomic DNA following the presented protocol (S1 Supplementary Protocol) and applied size selection using the Short Read Eliminator (SRE) kit (Circulomics), an easy-to-use method that does not require dedicated devices which is based on a length-dependent precipitation of nucleic acids driven by polyvinylpyrrolidone crowding. Large amounts of small DNA fragments can be detrimental for long-read Nanopore sequencing (14), not only because the subsequent reads are short, but also because these molecules can outcompete the longer ones, both for adapter ligation and pore usage, thus yielding suboptimal results.

The size distribution of the extracted DNA was assessed by PFGE and Nanopore sequencing, with and without size-selection for HMW DNA. Samples were migrated in a pulse field, stained by ethidium bromide and imaged with UV light (Fig. 1a). The DNA molecules extracted without size selection migrated as a large smear spread between approximately 1.5 and 150 kb. After size selection with the SRE kit, the upper part of the distribution remained unchanged while the low-molecular-weight fragments (< 10 kb) were visibly reduced. We made a similar observation after electrophoresis and staining of the samples in a 0.3% agarose gel (Fig. 1b).

**Figure 1.**
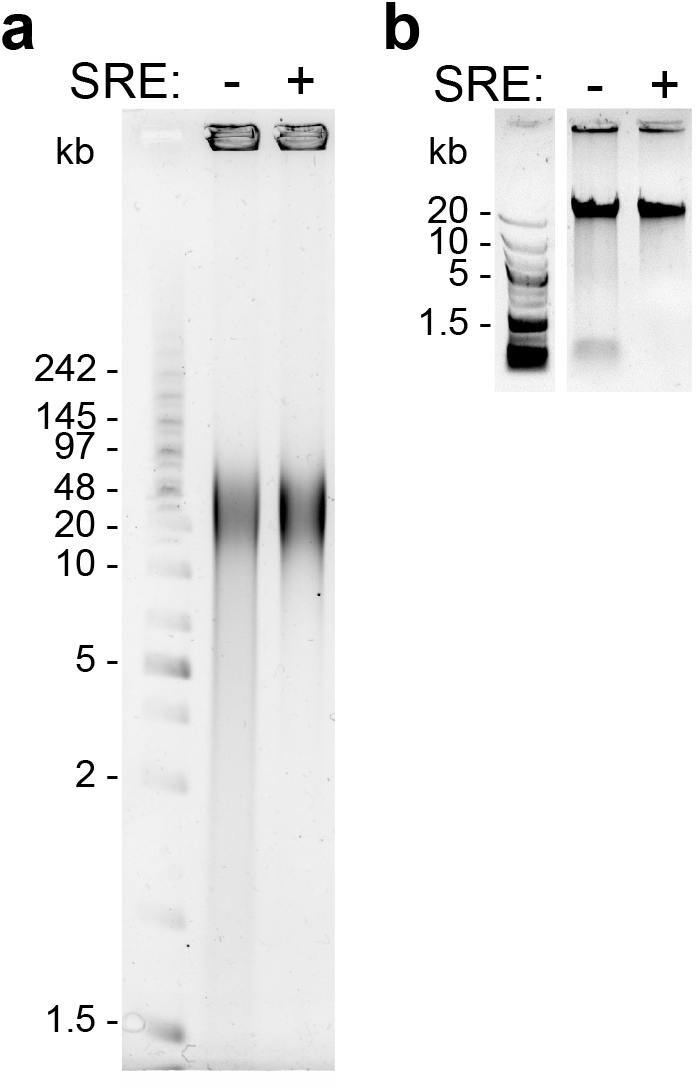
Visualization of extracted genomic DNA size distributions. (a) PFGE using 0.5 μg of DNA prepared with (+) or without (-) SRE size-selection, embedded in 30 μl of 0.5% low-melting agarose plugs, migrated in a 1% SeaKem GTG agarose (Lonza) gel. The ladder is a mix of PFG mid-range (N0342S, NEB) and GeneRuler 1 kb Plus (SM1331, ThermoFischer). Electrophoresis conditions: 0.5X TBE (Tris Borate EDTA) buffer, 6 V.cm^-1^, 120° angle, for 11h, switching time ramp from 1 to 60 seconds. Gel stained in ethidium bromide and imaged with UV. (b) Standard gel electrophoresis (0.3% agarose) of the indicated samples. GeneRuler 1 kb Plus (SM1331, ThermoFischer) is used as the ladder.

Size-selection of DNA fragments before preparation of libraries for Nanopore sequencing reproducibly led to a substantially decreased number of shorter molecules and an enrichment of longer ones (Fig. 2a-c), without negatively affecting read quality (Fig. 2d) and with no effect on genome-wide sequencing depth (S3 Supplementary Figure). Size-selection doubled the mean read length, increased the N50 from 12 kb to 17 kb (20 kb in two other experiments), with reads in the top decile being longer than 21 kb (26 kb and 27 kb in two other experiments) (S2 Table). The longest molecules we sequenced were over 100 kb, which are instrumental for genome assemblies.

**Figure 2.**
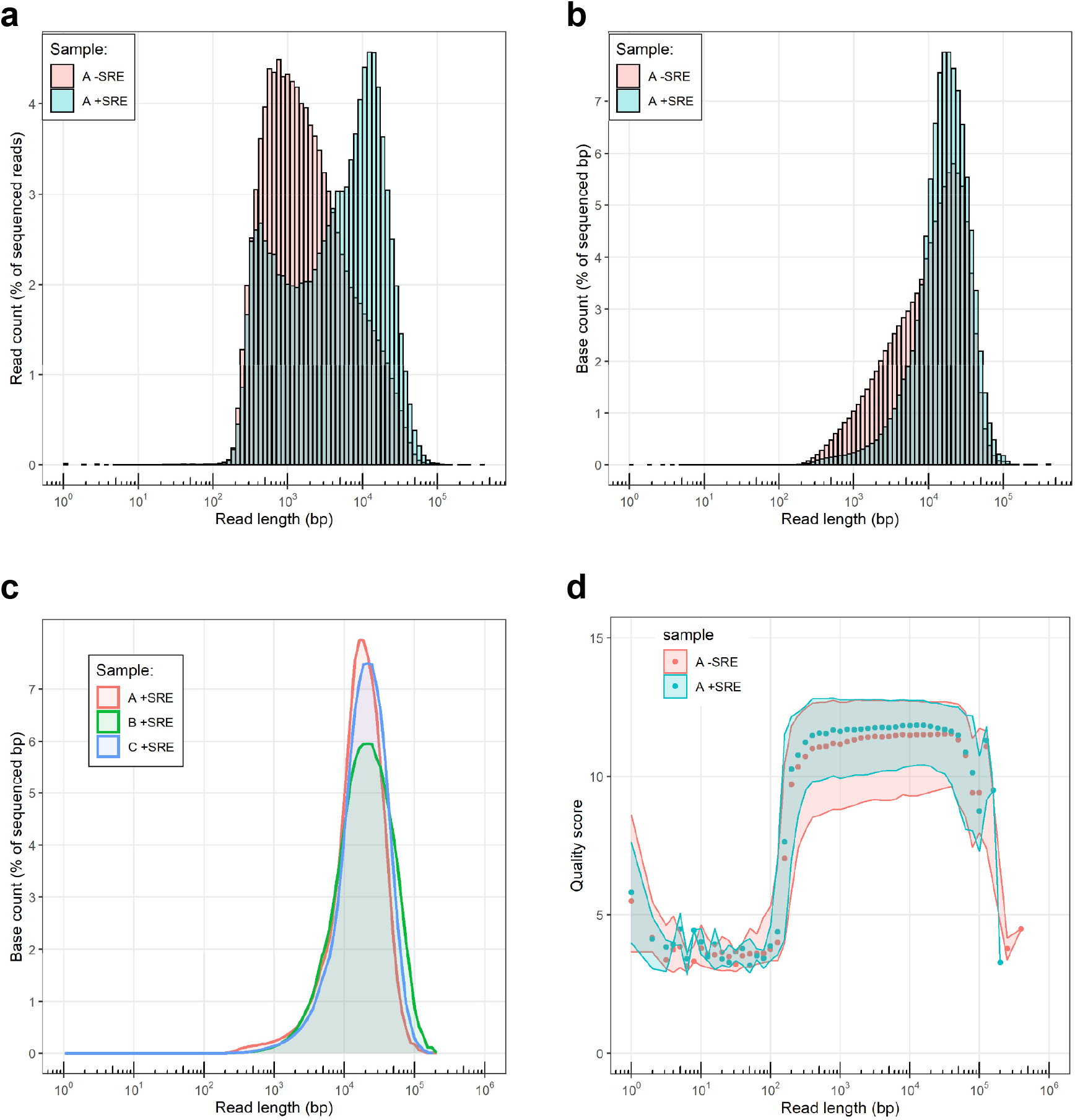
Distributions of read length in Nanopore-sequenced datasets. (a-b) Count percentage of (a) reads and of (b) bases as a function of read length obtained from genomic DNA of *C. reinhardtii* (experiment “A”, see Table S2) with or without size selection (+SRE and -SRE). (c) Count of bases after size-selection (+SRE) as a function of read length obtained from three different samples (see S2 Table and S4 Supplementary Figure). (d) Quality score for individual reads, grouped into bins of 0.1 log unit for samples “A-SRE” and “A+SRE”. The shaded areas represent the values between the 1^st^ and 3^rd^ quartiles.

## Acknowledgements

We thank Samuel O’Donnell for his help in the initial development of this protocol.

## Supporting information

### S1 Protocol

Step-by-step protocol, also available on protocols.io: dx.doi.org/10.17504/protocols.io.8epv59j9jg1b/v2

**S2 Table.** Summary statistics for 6 DNA preparations and sequencing experiments. Major limiting outputs are shown in red.

| Sample | CLIP library<sup>a</sup> | DNA extraction | Library preparation | Read count | Base count (Gb) | Read length quantiles (kb) |  |  | Median read length weighted by base count ("N50") (kb) | Top 3 longest reads<sup>b</sup> (kb) |  |  |
| --- | --- | --- | --- | --- | --- | --- | --- | --- | --- | --- | --- | --- |
|  |  |  |  |  |  | Median | Q90 | Q99 |  |  |  |  |
| A+SRE | LMJ.RY0402.077111 | This protocol | LSK109 +barcodes | 232 076 | 2,0 | 5,0 | 21 | 44 | 17 | 142 | 121 | 120 |
| A-SRE |  | This protocol (except SRE) | LSK109 | 850 066 | 3,6 | 1,5 | 12 | 34 | 12 | 127 | 117 | 112 |
| B+SRE | CC-4533 (cMJ030) | This protocol | LSK109 | 280 614 | 3,1 | 6,7 | 26 | 64 | 20 | 213 | 201 | 191 |
| B-SRE |  | Monarch HMW DNA kit<sup>c</sup> | LSK109 | 177 794 | 1,1 | 2,5 | 15 | 53 | 15 | 199 | 177 | 175 |
| C+SRE | LMJ.RY0402.209904 | This protocol | LSK110 | 344 223 | 3,9 | 8,0 | 27 | 54 | 20 | 175 | 166 | 161 |
| C-SRE |  | DNeasy Plant/Tip100<sup>d</sup> | LSK109 | 81 078 | 0,6 | 3,0 | 22 | 54 | 21 | 121 | 119 | 108 |

<sup>a</sup> https://www.chlamylibrary.org and reference (15).
<sup>b</sup> with quality > 7.
<sup>c</sup> as per manufacturer's protocol (Monarch® HMW DNA Extraction Kit for Tissue Cat. no. T3060L, New England Biolabs).
<sup>d</sup> cell lysis using DNeasy Maxi Plant (Cat. no. 68163, Qiagen) as in (16) and purification using Genomic-tip 100/G (Cat. no. 10243, Qiagen), then AMPure beads (Cat. no. A63880, Beckman Coulter).

**S3 Supplementary Figure.**
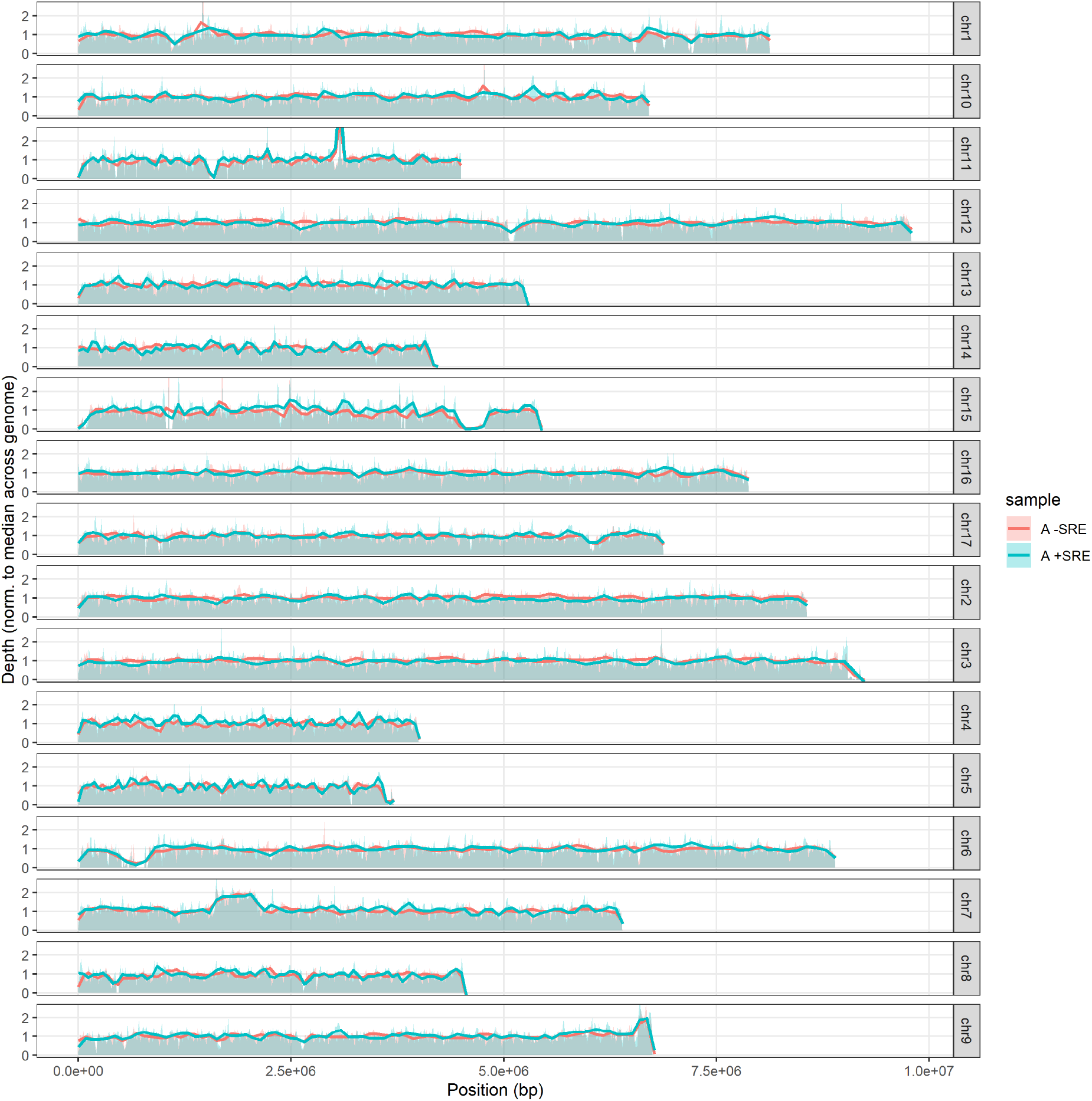
Sequencing depth normalized to the median across the whole genome of the sequencing reads for all chromosomes, using DNA obtained with (+) or without (-) SRE size selection.

**S4 Supplementary Figure.**
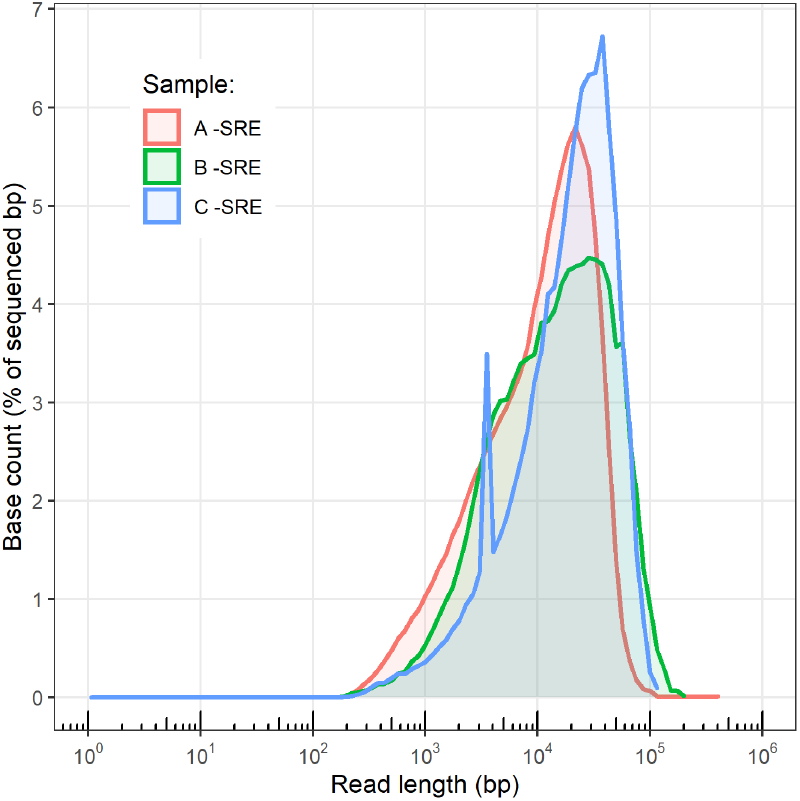
Count percentage of bases as a function of read length with alternative sample preparations without size selection (-SRE). See Table S2 for details. Sample C was sequenced in the presence of control DNA (“DNA CS” from Oxford Nanopore sequencing), which peaked at 3 kb.

## Notes

https://www.protocols.io/view/extraction-and-selection-of-high-molecular-weight-b9pir5ke

### Competing Interest Statement

The authors have declared no competing interest.

